# Genes and genome-resolved metagenomics reveal the microbial functional make up of Amazon peatlands under geochemical gradients

**DOI:** 10.1101/2022.12.19.521090

**Authors:** Michael J. Pavia, Damien Finn, Franco Macedo-Tafur, Rodil Tello-Espinoza, Christa Penaccio, Nicholas Bouskill, Hinsby Cadillo-Quiroz

## Abstract

The Pastaza-Marañón Foreland Basin (PMFB) holds the most extensive tropical peatland area in South America. PMFB peatlands store 7.07 Gt of organic carbon interacting with multiple microbial heterotrophic, methanogenic, and other oxic/anoxic respirations. Little is understood about the contribution of distinct microbial community members inhabiting tropical peatlands. Here, we studied the metagenomes of three geochemically distinct peatlands spanning minerotrophic, mixed, and ombrotrophic conditions. Using gene- and genome-centric approaches, we evaluate the functional potential of the underlying microbial communities. Abundance analyses shows significant differences in C, N, P, and S acquisition genes. Further, community interactions mediated by Toxin-antitoxin and CRISPR-Cas systems were enriched in oligotrophic soils, suggesting that non-metabolic interactions may exert additional controls in low nutrient environments. Similarly, we reconstructed 519 metagenome-assembled genomes spanning 28 phyla. Our analyses detail key differences across the nutrient gradient in the predicted microbial populations involved in degradation of organic matter, and the cycling of N and S. Notably, we observed differences in the nitrogen oxide (NO) reduction strategies between sites with high and low N_2_O fluxes and found phyla putatively capable of both NO and sulfate reduction. Our findings detail how gene abundances and microbial populations are influenced by geochemical differences in tropical peatlands.

## Introduction

Tropical peatlands are globally important carbon reservoirs that span across continents[1], but are vulnerable to deterioration from anthropogenic influences, such as land conversion for agriculture and the changing climate[2]. Two critical roles of tropical peatlands are their capacity to store soil organic matter (SOM) and the emission of greenhouse gases (GHG) when SOM is degraded[3, 4]. In the western Amazon, the Pastaza-Marañón Foreland Basin (PMFB) is the largest known area of tropical peatlands in South America, representing close to 120,000 km^2^ from recent estimates[4–6]. In effect, the PMFB stores ∼7.07 Gt of carbon in soils and its decomposition contributes an estimated 31.6 – 41.1 Tg methane (CH_4_) year^−1^ [5]. However, the carbon storage capacity of the PMFB is predicted to decrease within this century as a response to climate change scenarios [7–9]. The decline in carbon storage capacity of the PMFB will result in the further release of CO_2_ and CH_4_ to the atmosphere through microbial processes, representing a positive feedback to climate change. Understanding the processes and microbes that affect SOM decomposition in Amazon peatlands is needed to better evaluate and predict soil carbon stability and GHG emissions in the region.

Heterogeneity in interannual rainfall and/or riverine flood, along with each site’s geomorphology, has led to strong site-to-site variability in the concentration and availability of biologically important elements (Ca, Mg, K, Na, Fe, P, SO_4_ and others) across peatland soils in the PMFB[10]. Thus, minerotrophic, mixed, and ombrotrophic soil conditions are common in the region[11–13]. Soil geochemistry plays an important role in governing microbial community structure at both taxonomic and functional gene scales[14–16]. Variations in microbial community composition are especially noticeable when comparing minerotrophic sites— characterized by higher concentrations of minerals and closer to neutral pH—*versus* rainfall-fed ombrotrophic sites with far lower concentrations of minerals and a more acidic pH. Despite a scarcity of available data, preliminary observations indicate that tropical minerotrophic sites are predominately inhabited by *Alphaproteobacteria, Gammaproteobacteria*, and hydrogen metabolizing *Archaea*[13, 17]. By contrast, *Acidobacteria* and *Verrucomicrobia* are the dominant community members in ombrotrophic tropical peatlands[13, 18]. In particular, mineral concentration and pH are key environmental factors which contribute to variations in the emergent microbial community composition across the PMFB[13, 19]. Metabolic capabilities and survival strategies of microbial populations in ombrotrophic to minerotrophic soil can directly influence the rate at which stored SOM is converted into GHGs. Identifying the community composition and analyzing its variation within their geochemical context has been successful in initial predictions of the functional potential found in peat soils[13, 19–22]. However, this leaves a gap in our knowledge of the genetic or putative traits contained within these microbial populations driving biogeochemical cycles in the soil.

Understanding how microbial communities cope with environmental variation requires incorporating both a community scale view of functional gene make up, as well as gene combinations within members of the community. Metagenomic approaches can describe the taxonomic and functional potential of microbial populations within highly complex and diverse communities. Of the few available studies from tropical peatlands, important findings have proposed a link in soil properties to the nitrogen cycling community [22], as well as identified potential key phyla involved in the degradation of plant polysaccharides[21]. Furthermore, binning of sequences from metagenomes have facilitated the recovery of poorly characterized taxonomic clades that are not amenable to cultivation, and identified key microbial populations involved in degradation of SOM[23]. While nutrient availability drives community member functions, cellular traits associated with homeostasis and survival strategies have also been shown to be regulated by soil properties[24]. Many of these functional traits are evolutionarily selected for in response to environmental conditions. In general, metagenomic studies of peatlands have been concentrated in the northern hemisphere, but have provided valuable insights into key functions and microbial populations involved in the biogeochemical cycles occurring in northern peat soils[17, 23, 25]. Metagenomic studies in tropical peatlands are rare and, to our knowledge, there is no published metagenomic study for peatlands found in the Amazon. To date, only two studies have looked at the microbial community member abundance differences across the PMFB nutrient gradient using the 16S rRNA marker gene, however we lack an understanding of the functional gene landscape and how these are partitioned among prokaryotic populations. In this metagenome analysis, we hypothesize that (1) the functional gene landscape will reveal that variations in carbon processing and nutrient cycling pathways are influenced by geochemical differences, and (2) that adaptations to the underlying geochemistry have led to functional specialization of microbial populations across sites when compared to the larger ecosystem.

## Materials and methods

### Study sites, Sample Collection, DNA Sequencing and Metagenomic Library Processing

We studied the soil microbial communities from three peatlands within the PMFB: Buena Vista (BVA), Quistococha (QUI), and San Jorge (SJO). These study sites capture a broad range of heterogeneity of these peatlands, which include gradients in geochemistry, pH, hydrology, vegetation, GHG flux and microbial composition as detailed elsewhere[10, 13, 19, 26] and summarized in Supplementary Table 1. Soil core samples were collected at two depths (0-10 and 10-20 cm) as previously described in Finn *et al*., (2020) based on the younger nature of organic soils and higher microbial activity in surface layers. Briefly, sampling occurred in July 2015, August 2015, October 2015, January 2016, and February 2016 (Supplementary File 1) coinciding with expeditions at the peak of dry season and early wet season before regional flooding as periods where the microbial activity and composition is likely most distincy and thus sought to be captured. Triplicate 0.5 grams of soil were placed in sterile tubes with MoBio Lifeguard stabilization solution (MoBio, United States). Samples were refrigerated for 24-48 hours during transport to facilities, and stored at -80° C until DNA was extracted according to the protocol outlined in Lim, *et al*., (2016)[27]. Metagenome libraries were prepared using 40-200 ng of DNA and were sheared to ∼450 bp using a Covaris LE220 focused-ultrasonicator. The sheared DNA fragments were size selected by double-SPRI and then the selected fragments were end-repaired, A-tailed, and ligated with Illumina compatible sequencing adaptors from IDT containing a unique molecular index barcode for each sample library, and libraries sequenced on Illumina Hi Seq 2500 2 x 151 bp technologies at the Joint Genome Institute (JGI). Contamination, read trimming, and quality score filtering were also carried out in accordance with JGI IMG protocol[28], and average coverage and sequence diversity of the metagenomes were estimated using Nonpareil (v3.2) (Supplementary File 1) [29].

### Determination of community level functional differences

To determine the relative abundance of functional genes, metagenomes were assembled using MEGAHIT (v1.1.3) with default parameters and the following k-mer list options “23,43,63,83,103,123” [30]. Post-assembly processing included quality assessment using QUAST (v3.0) [31], and open reading frame (ORF) identification using Prodigal (v 2.6.3)[32]. Interpretation of gene differences between sites was carried out by annotating ORFs against the KEGG Orthology database[33]. Annotations were filtered by an E value cutoff of less than 1E-7, length greater than twenty-five amino acids, and greater than 30% amino acid identity[34, 35]. The relative abundance of functional genes was quantified using a count table generated from the number of ORFs assigned to unique KO terms and used as a proxy for gene abundance within the microbial communities. To investigate community level differences in gene abundance we carried out a distance-based reduction analysis on gene counts using the Bray-Curtis dissimilarity index with the capscale function in R’s vegan package (v2.6-2)[36], and significance was tested using PERMANOVA with the ADONIS function in R’s vegan package. Further, hierarchical clustering of functional count data was performed with the stats package in R using the ward.D2 method. To identify the genes driving the dissimilarity between sites we implemented the DESeq2 package (v1.31)[37]. Subsets of significant genes, based on proposed cellular function, were then tested for significance using an ANOVA. All P values of < 0.05 were considered significant.

### MAG recovery and downstream analysis

Accurate binning of metagenome assembled genomes (MAGs) is difficult in soil because of the inherent community complexity[13, 21]. In this study, we employed a co-assembly approach to increase the probability of recovering high quality MAGs, by increasing availability of reads from low abundant populations. Quality controlled and trimmed metagenomes were pooled based on site and sampling year (Supplementary File 1). Metagenomes collected in February 2016 from BVA were co-assembled by sampling site instead of sampling year to increase BVA MAG representation. Assembly of merged metagenomes was done using MEGAHIT with default settings in addition to the preset meta-large [30]. Contigs in which 1 x coverage was achieved on 90% of their length were kept for downstream binning.

Contig binning was carried out with a minimum length of 1500 bp using the four binning programs MaxBin (v2.2.5) [38] (both bacterial and archaeal single copy genes (scg) options), MetaBAT2 (v2.12.1) [39], CONCOCT (v0.4) [40], and BMC3C[41]. The resulting bins were consolidated using DAS Tool (v1.1)[42], and then further curated using a combination of RefineM (v0.1.1)[43] and Anvi’o (v6.1) [44]. Curated MAGs were assessed for completeness and redundancy using CheckM [45]. MAGs whose quality were >50% complete and <10% redundant were classified using GTDB-Tk against the Genome Taxonomy Database release 07-RS207 [46], and a phylogenetic tree was generated using GToTree (v1.6.37) [47] and visualized using ggtree [48]. Relative abundance of MAGs was determined first by mapping individual metagenomes to co-assembled contigs using Bowtie2[49]. Then a JGI script provided in MetaBat[50] was used to calculate contig coverage and normalized by MAG size. ORFs in MAGs were identified using Prodigal[32], and annotated using a combination of GhostKoala[51] and MagicLamp[52]. Pathways for cycling of organic and inorganic compounds encoded within MAGs were identified using automated methods based on the presence or absence of pathway genes with a threshold determined by the completeness of the MAG. Finally, alpha diversity for MAGs predicted to be involved in polysaccharide degradation was calculated using the R vegan package[53].

## Results

### Community diversity

Retrieval of metabolic potential and population dynamics patterns were targeted from 24 metagenomes sequenced from Buena Vista (BVA), Quistococha (QUI), and San Jorge (SJO) peatlands that represent a nutrient gradient, from high to low, respectively, in the PMFB[13, 19]. A detailed description of the samples with corresponding geochemistry from the soils cores can be found in Finn *et al*., 2021 [13] and also summarized in Supplementary Table 1. Estimated community coverage and read sequence diversity, calculated by Nonpareil 3[29], varied as a function of nutrient status (Two-way ANOVA, *p* < 0.05) (Supplementary Figure 1). We detected a higher read sequence diversity (alpha diversity) and low community coverage associated with the nutrient-rich metagenomes (BVA), while lower sequence diversity and high coverage was achieved in nutrient-poor metagenomes (SJO) and intermediate values in the intermediate site (QUI), albeit nutrient poor-leaning.

**Figure 1.**
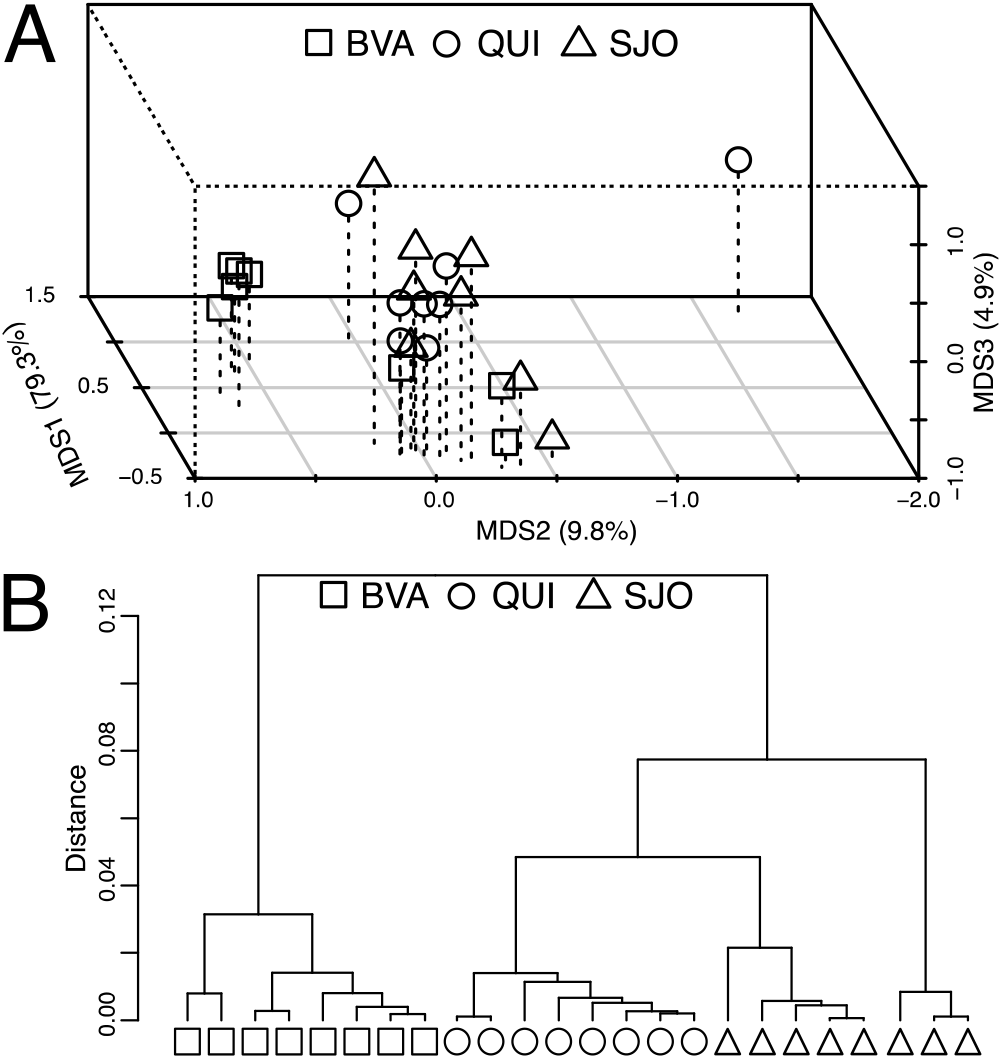
Clustering comparisons of amazon metagenomes. (A) Bray-Curtis dissimilarity cluster analysis of KEGG derived functional annotations (community level functional abundance) of genes recovered in metagenomes from Amazon peatlands in study. (B) Hierarchical clustering of functional abundance variance same dataset in study. ANOSIM : p = 0.002

### Community level gene abundance differences across the mineral nutrient gradient

Gene frequency, based on Bray-Curtis dissimilarities of KEGG-derived gene annotations, show distinct separation between peatland ecosystems (ANOSIM, *p* < 0.05) (Fig. 1A). Hierarchical clustering was consistent with Bray-Curtis dissimilarity, but overall marginal variation was observed in a subset of SJO samples (Fig. 1B). Of the 10 968 annotated genes detected across all sites, 3 502 genes were identified by DESeq2 as differentially abundant, based on Benjamini and Hochberg p-value adjustment, between metagenomes from the three peatlands (Fig. 2, Supplementary File 2). Within differentially abundant genes SOM processing, inorganic nutrient cycling, and genes potentially involved in ecological interactions were found to have distinct patterns between BVA, QUI, and SJO. Overall, genes involved in degradation of oligosaccharides and aromatic hydrocarbons were enriched within nutrient-poor sites QUI and SJO. However, downstream processing via pentose phosphate pathway, benzoate, and furfural degradation were only enriched within the most oligotrophic site, SJO. Interestingly, genes associated with methanogenesis were enriched within the nutrient rich BVA; yet, a gene involved in aerobic methanotrophy, *mmoX*, was enriched in metagenomes from QUI and SJO.

**Figure 2.**
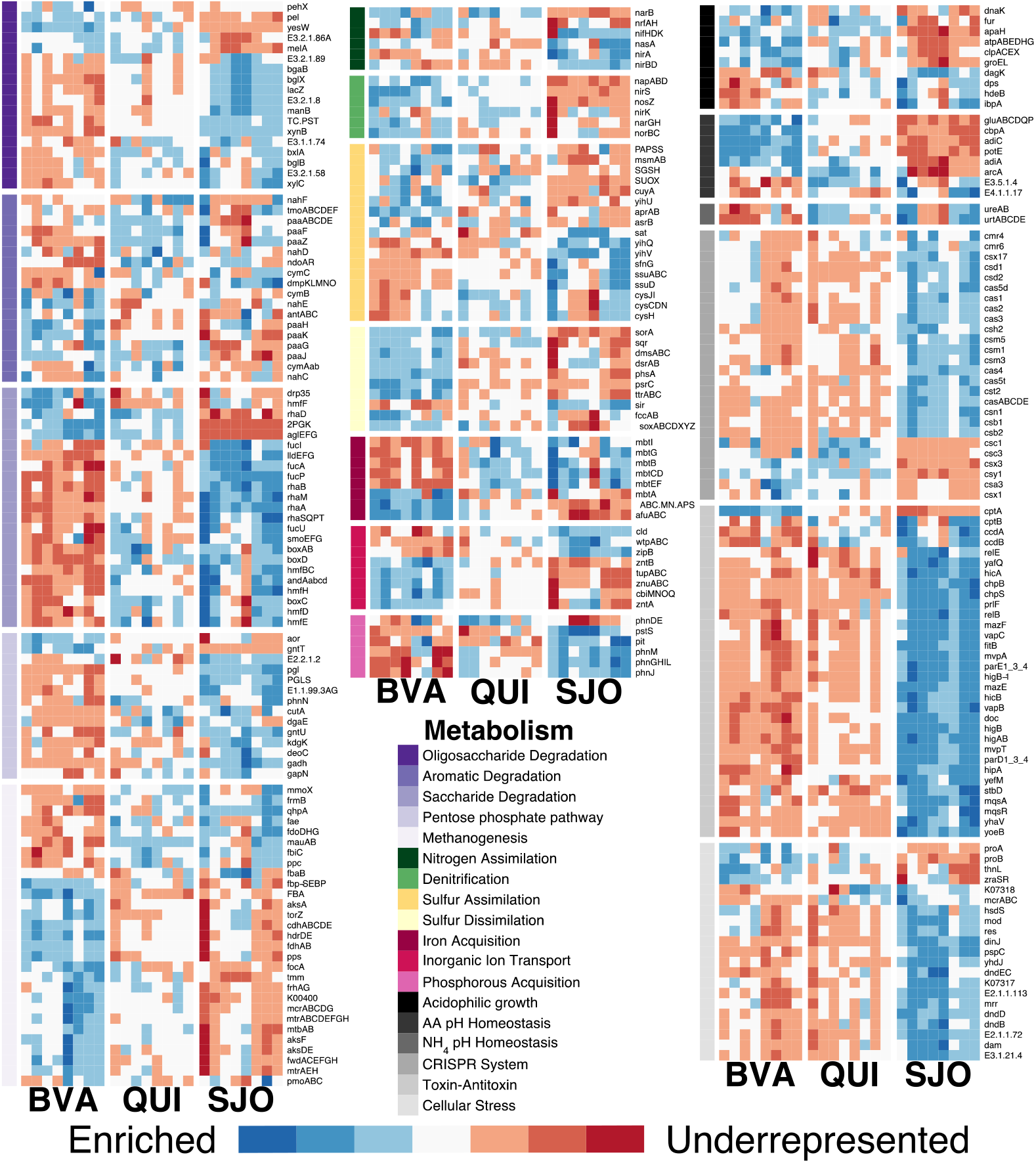
Heatmap of log-fold differences in abundance of selected genes identified by DESeq2 as being significantly different across the PMFB nutrient gradient. Color bars on the left indicate the subsystems and labels on the right are the gene/gene complex names. For genes associated with a protein complex the log fold difference shown is the average of all subunits. Color code is based on the effect size estimate and scaled by row. * Denotes a p<0.05 for a two-way ANOVA on the genes within the subset between the three sites (BVA, QUI, and SJO). We note that pmoABC was not identified as being significantly different but is plotted because of its importance in methanotrophic metabolism.

Nitrogen and sulfur transformations for both assimilatory and dissimilatory pathways were differentially abundant between the three peatlands. Genes involved in nitrogen assimilation and fixation were abundant in SJO, while in BVA and QUI metagenomes, *narB* was more abundant while *nrfAH* is more abundant in BVA only; in contrast denitrification genes were significantly enriched within BVA and at low abundance in QUI and SJO. Sulfur assimilation and dissimilation pathways were enriched in SJO and BVA, respectively. Siderophores, iron-chelating molecules used for iron acquisition, were enriched in the nutrient-poor sites QUI and SJO, while direct iron ion uptake was enriched in BVA. Genes involved in cycling of inorganic compounds show a unique pattern associated with the nutrient gradient across the PMFB.

Adaptations to soil acidity and community interactions also differed between sites. Amino-acid dependent pH survival strategies were highly enriched within BVA, which contrasted with the enrichment of ammonia generation genes as a potential pH survival strategy in SJO. Genes for both types of pH survival strategies were enriched in QUI. Most notably genes that can be grouped into prokaryotic survival strategies, such as those involved in CRISPR-Cas system, Toxin-antitoxin systems (TA systems), and DNA repair, were enriched in the oligotrophic SJO metagenomes. TA systems are commonly utilized by microbes for stress response and/or dormancy[54]. Thirty-two genes, representing eleven complete TA systems, were significantly enriched in SJO with the most common functions identified as regulation of macromolecule biosynthesis and plasmid population. Overall, gene abundance differences across the PMFB nutrient gradient were greatest between the nutrient-rich site BVA and nutrient-poor site SJO, with QUI sharing more common enrichment of functions with SJO.

### MAG-resolved composition and their functional potential across the nutrient gradient

To better understand the genome resolved make-up of microbial clades and their potential ecological roles across the PMFB, all metagenomes were co-assembled by sampling date with the exception of BVA metagenomes from February 2016 which were co-assembled by sampling site to improve recovery of quality MAGs (Supplementary File 1). On average, contigs used for binning recruited 59% (BVA), 67% (QUI), and 79% (SJO) of reads from individual metagenomes. From all co-assemblies, a total of 2 225 MAGs were recovered, with an average of 163, 266, and 348 MAGs, from the three sites, BVA, QUI, and SJO, respectively. Co-assembly and binning statistics for each site are summarized in Supplementary File 1. We recovered a total of 519 MAGs, with >50% completeness and <10% redundancy from the nine co-assemblies representing both bacterial (398 MAGs) and archaeal lineages (121 MAGs) (Fig. 3, Supplementary File 3). The recovered MAGs span 27 phyla, including *Bacteria* belonging to *Acidobacteriota* (87), *Actinobacteriota* (50), and *Nitrospirota* (35), and *Archaea* from *Halobacteriota* (24) and *Thermoplasmatota* (23). MAGs from poorly characterized phyla *Dromibacteria, Eisenbacteria*, and *Vampirovibrionia* were also recovered. High to medium quality MAGs recovered from BVA (92) were greatly underrepresented when compared to the those from QUI (222) and SJO (205) (Supplementary File 1).

**Figure 3.**
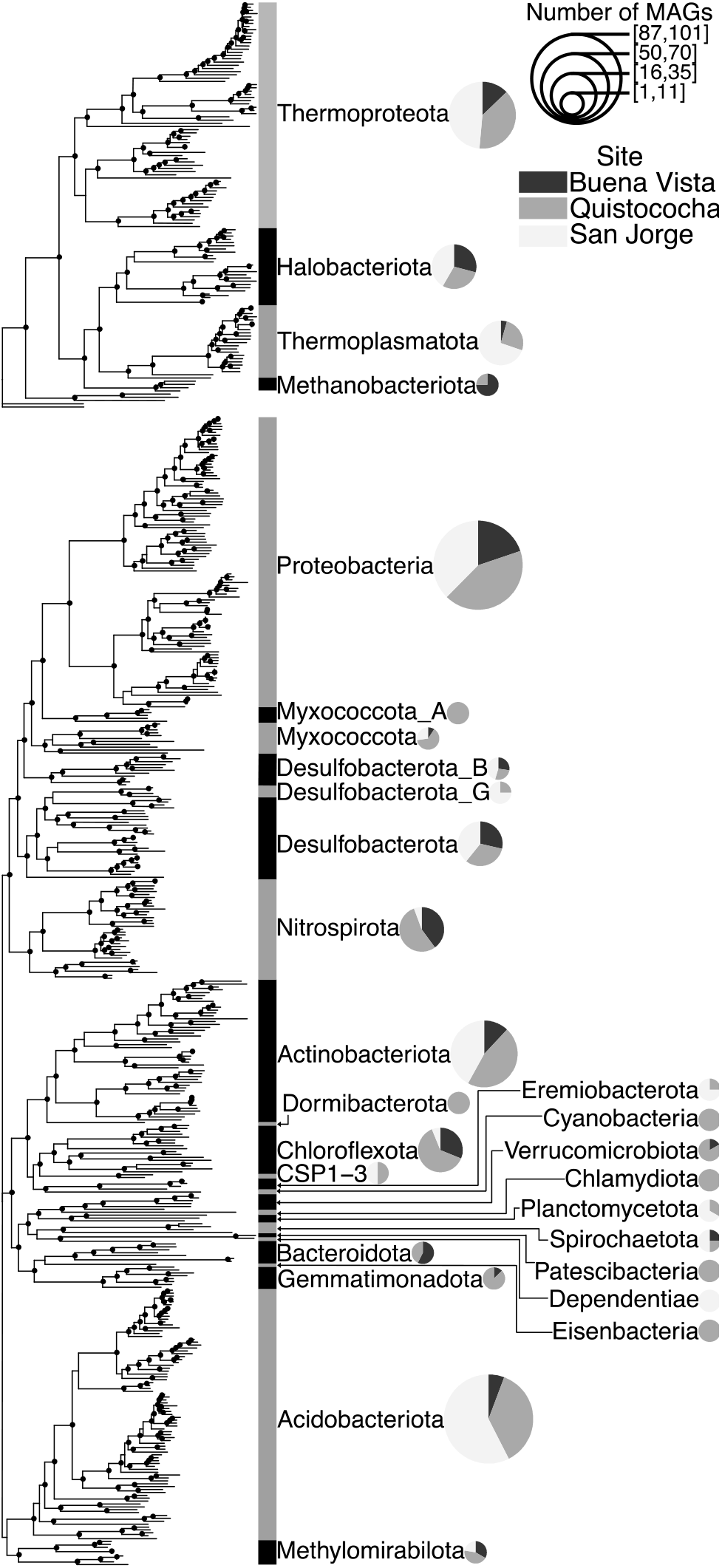
Phylogenetic placement of bacterial and archaeal MAGs recovered from the PMFB metagenomes. Filled circles at branches represent a bootstrap score greater than 70%. MAG based community profile at phylum level. Pie charts by Phylum represent the proportion of MAGs from each site, which is scaled in size to represent the number of MAGs recovered in this study.

Reconstructing the metabolic potential and average abundance of MAGs within each site, allowed for the evaluation of key metabolic pathways that lead to or affect the production of greenhouse gasses from the PMFB (Fig. 4). Breakdown of polysaccharides into monomers for primary and secondary fermentation were predominantly associated with MAGs, across all three sites, identified as *Acidobacteriae* (Fig. 4A). The *Acidobacteriae* appear to be the primary bacterial degraders of polysaccharides in SJO, while BVA and QUI populations potentially responsible for polysaccharide degradation were higher in recovered MAG diversity, associated primarily to *Gemmatimonadetes* and *Thermoplasmata*, in addition to the *Acidobacteriae*. Genetic potential for cutin degradation were found in MAGs from QUI and SJO, although were only found in *Actinomycetia* MAGs. Unique to BVA, *Actionmycetia* MAGs were found to have the genetic potential for the degradation of xylose (both isomerase and oxidoreductase pathways) and cellulose. Further, degradation of saccharides, via primary and secondary fermentation, were higher in recovered MAG diversity in BVA and QUI when compared to SJO (BVA: 3.19, QUI: 3.35, and SJO: 2.72). Irrespective of site, but prominent in QUI and SJO, MAGs classified as *Acidobacteriae*, were found to contain the genetic potential for four out of five sugar fermentation pathways. Fermentation products (CO_2_, acetate, methylated single carbon compounds and H_2_) are precursory products for anaerobic carbon transformation to CH_4_ by methanogenic archaea. Fixation of CO_2_ can offset availability and reduce fuel for hydrogenotrophic methanogens. Within QUI and SJO, the genetic potential for CO_2_ fixation was found in *Acidobateriae* and *Bathyarchaeia* MAGs, while *Gammaproteobacteria* were found to be putative CO_2_ fixers within BVA.

**Figure 4.**
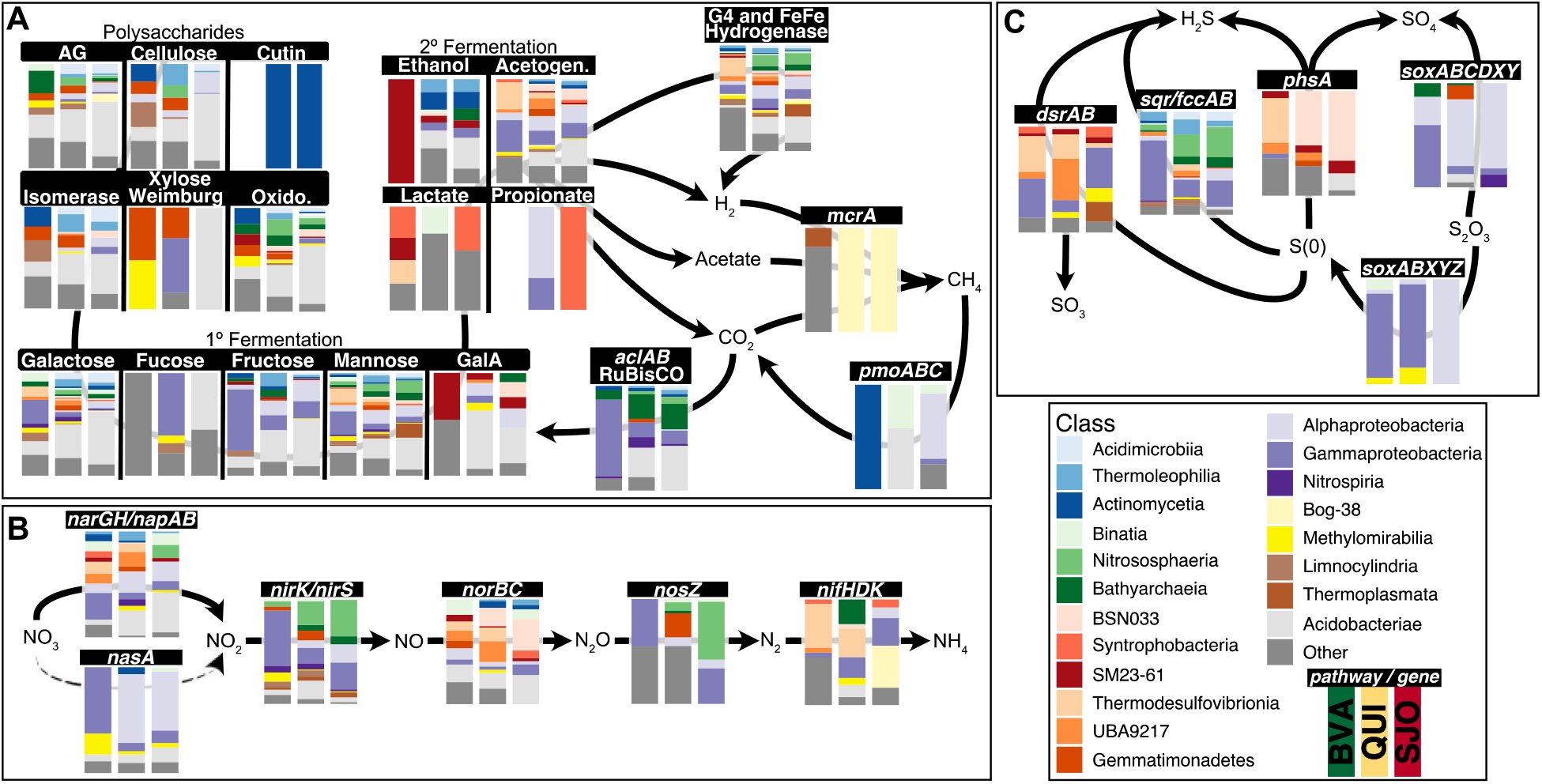
Proportion of MAG relative abundance across BVA, QUI, and SJO sites capable of (A) carbon degradation and cycling of (B) Nitrogen and (C) Sulfur. Solid arrows indicate degradative or dissimilatory transformations, and dashed arrows indicated assimilatory transformations. The presence of a carbon pathway was determined based on percentage of genes in pathway detected normalized by MAG completeness (Supplementary File 4). Abbreviations used in A are as follows, AG : Arabinogalactan, Oxido : Oxidoreductase, Acetogen. : Acetogenesis, and GalA : Galacturonic Acid. For B and C, genetic potential of multigene complexes is based on the presence of at least the catalytic subunit (Supplementary File 4). MAG classifications achieved different levels (Supplementary File 3) but is summarized here at the class level as detailed in legend.

The genetic potential for conversion of secondary fermentation products to CH_4_ (*mcrA*), was found exclusively in *Halobacterota* MAGs, except for one MAG classified to Thermoplasmata from BVA (Fig. 4A). Both acetoclastic and hydrogenotrophic MAGs were recovered from the nutrient rich BVA, *Methanotrichaceae* and *Methanocellaceae* (contained in the “other” category of Fig. 4A), respectively. Conversely, only hydrogenotrophic methanogens, classified as *Methanomicrobia* (Bog-38 in the GTDB database), were recovered from QUI and SJO. The genetic potential to oxidize CH_4_ into biomass (*pmoABC*) was found to be contained within different phyla across the three sites (Fig 4A). In BVA, our sampling was able to recover one population with the genetic potential capable of CH_4_ oxidation belonging to the *Actinomycetia* class (family *Kineosporiaceae*). Given the unusual finding for potential methanotrophy in this phylum, we inspected the MAG and found no evidence for contamination or ambiguous reads assignments, and instead also found genetic potential for nitrate/nitrite reduction. It remains to be elucidated whether this population is capable of canonical aerobic, or nitrate/nitrite-dependent anerobic methanotrophy, or none. MAGs classified to *Candidatus* Binatia with *pmoABC* were found in both QUI and SJO, and the putative methanotrophic capacity for this group has also been postulated elsewhere[55]. Overall, many steps in the carbon degradation cascade have potential to be carried out by multiple microbial populations across the nutrient gradient, with the most distinct differences occurring in highly conserved genes, *mcrA* and *pmoABC*.

Microbial transformations of sulfur and nitrogen compounds can affect N_2_O, CH_4_, and CO_2_ fluxes from soils. Both high sulfate reduction rates and high concentrations of N_2_O have been shown to inhibit methanogenesis[56, 57]. Across the nutrient gradient MAGs with the genetic potential for sulfate reduction (*dsrAB*) were distinct between the three sites (Figure 4C). *UBA9217* MAGs, abundant at QUI, were found to be capable of sulfate reduction which contrasts with the abundant *Gammaproteobacteria* MAGs recovered in BVA and SJO. Furthermore, the genetic potential to produce SO_4_ via disproportionation of S° (*phsA*) was found in abundant *Thermodesulfovibrionia* MAGs from BVA, but abundant BSN033 MAGs in QUI and SJO. Alternatively, thiosulfate oxidation via the complete SOX pathway was found among *Alphaproteobacteria* MAGs in QUI and SJO, while *Gammaproteobacteria* were abundant within BVA.

Denitrification is responsible for removal of nitrogenous compounds from the soil as either N_2_O or N2. Overall, the genetic potential to transform NO_3_ to N2 was predominantly found within abundant *Gammaproteobacteria* in BVA or, abundant *Alphaproteobacteria* in QUI (Fig. 4B). In SJO, archaeal *Nitrososphaeria* MAGs were found to hold *narGH* genes suggesting some members could participate in early steps of denitrification, however the sporadic presence of *NirK* has been suggested to assist in the detoxification of ammonia oxidation [58] rather than a denitrification role in one order of this class. Contrastingly, abundant *Thermodesulfovibrionia* were found in both BVA and QUI as capable of nitrogen fixation (*nifHDK*), while the nitrogen fixing community in SJO is potentially dominated by the *Methanomicrobia* (Bog-38 in the GTDB database). High concentrations of N_2_O have been shown to experimentally inhibit methanogenesis in PMFB soils[19], thus we further explored the genetic potential for NO conversion to N_2_O. To evaluate putative NO reducers across the nutrient gradient, *norB* sequences were classified as either being used for energy conservation if electrons are accepted from the quinol pool (*qNor)* or detoxifiers if electrons are accepted from cytochrome c pool (*cNor)* (Figure 5). Many NO reducers were not capable of carrying out other steps in the canonical denitrification pathway regardless of site or taxonomic classification (Figure 5). In sites with higher N_2_O emissions (BVA and SJO) [13, 19] many of the NO reducing MAGs contained *qNor* (BVA:8, QUI:20, SJO:15). Within the NO reducing communities, regardless of *norB* clade, many were also capable of sulfate reduction. This was predominantly observed in NO detoxifying *Thermodesulfovibrionia* and UBA9217 MAGs in QUI. For BVA and SJO, *Desulfomonilia* and BSN033, respectively, were found to also be potentially NO and sulfate reducers, but only a few MAGs were identified as having this capability. Overall, NO reducers within the PMFB show differences in NO reduction strategy, where energy conservation is more prevalent within sites where higher N_2_O flux has been detected [13, 19].

**Figure 5.**
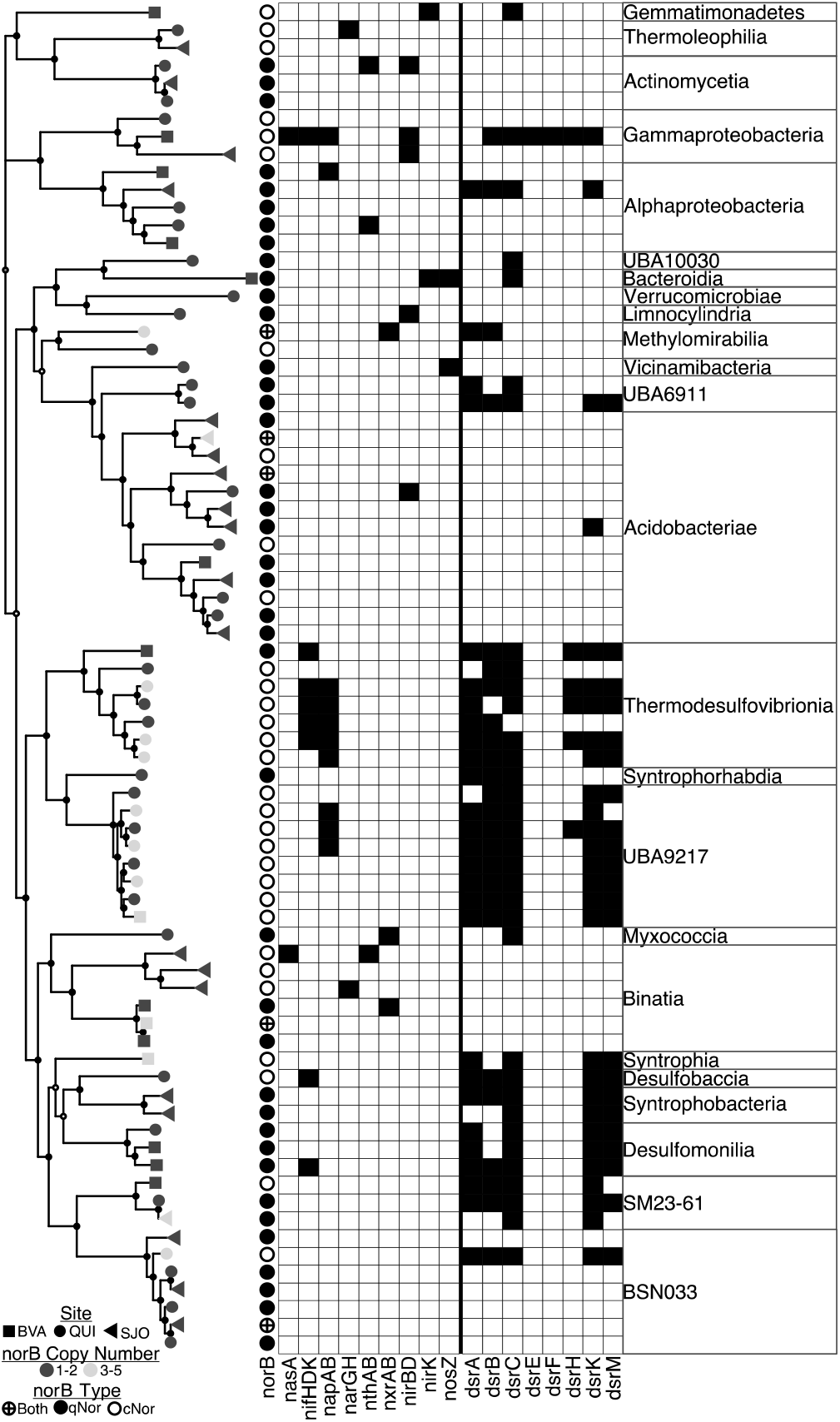
Nitrogen and sulfate cycling metabolic potential of potential NO reducing MAGs recovered from the PMFB. Filled circles at branches represent a bootstrap score greater than 70%. Shapes at tree tips represent the site from which that MAG was recovered from. Color of tree tip indicates the number of norB gene sequences found within that MAG. Circles with a cross show that both a qNor and cNor sequence was found in the MAG. Fill circle represents a qNor ORF and open circles represents a cNor ORF. Filled in squares represent the presence of the gene or gene complex for nitrogen and sulfur cycling.

## Discussion

### Community level functional differences reflect geochemical differences across sites

In this study, we took a metagenomic approach to understand differences in relative abundance of microbial functions across tropical peatlands conditions found within the PMFB. The soils of BVA, QUI, and SJO are characterized by geochemical differences with respect to pH, nutrient concentrations, and GHGs emissions (CH_4_, CO_2_, and N_2_O) [6, 11, 13]. In the selected PMFB study sites, higher nutrient concentrations (BVA) were associated with high metagenome read sequence diversity. These findings are consistent with other studies that have shown pH and nutrient concentrations are linked to both the demographic and functional diversity of the microbial community [20, 59]. For example, Finn et al. (2020) has shown that pH and nutrient concentrations are drivers of community composition, in particular the methanogens and methanotrophs of the PMFB. When detailing the dissimilarity of gene abundances in metagenomes across the three sites, we observed that their functional landscapes are significantly different, which suggest that their defined geochemical differences are also drivers of the functional potential and abundance. Further, clustering by gene abundance variance demonstrates that QUI and SJO metagenomes have more closely related genetic potential, in comparison to BVA. Thus, from a microbial functional perspective the oligotrophic (SJO) and intermediate-poor (QUI) sites can putatively (by gene content) share common pathways or mechanisms to address the challenges or opportunities in their peat soils. The lower nutrient conditions for SJO and QUI are driven by their dominant water source, rain-only in the former and mostly-rain with limited surface water in the latter, which are highly different to the riverine-flooded BVA[11].

When examining differential abundance of genes across the three sites, 36% of genes showed statistically significant differences in abundance. We focused on genes whose annotations are well documented, of which, most were found to be associated with carbon processing, inorganic nutrient cycling, and microbial survival strategies. These functions can directly or indirectly influence the various fluxes of GHGs measured across the PMFB [13, 19]. With regards to the carbon degradation cascade, we identified that SJO is notably enriched for genes involved in polysaccharide and aromatic hydrocarbon degrading pathways. The C/N in SJO is much higher when compared to BVA and QUI, but given the abundance of these pathways, it is likely that the majority of SOC is locked in recalcitrant compounds[11, 13]. This is supported by the higher rate of peat accumulation, higher humification index and lowest methanogenic flux observed in SJO[11, 13]. Conversely, genes involved in methanogenesis were shown to be enriched within BVA, coinciding with the higher CH_4_ flux records for this site [7, 11, 13]. This is further supported by a greater abundance of methanogens found within BVA in particular *Methanobacteriacea*[13].

In addition, many genes involved in the cycling of inorganic compounds, such as N, S, Fe, and P, were found to have distinct patterns between the three sites. Genes associated with denitrification and dissimilatory sulfur cycling, were on average more abundant in the nutrient-rich site of BVA. Many of these genes are involved in proton pumping (*narGH, norBC, nosZ, sqr, soxABCDXYZ, dsrAB*, and *phsA*), and enable niches powered by alternative terminal electron acceptors to be occupied[60]. We propose that the abundance of these genes are attributed to the minerotrophic conditions at this site where more potential niches can be available and occupied by diverse microbial populations[61, 62]. A complementary pattern was observed in the oligotrophic SJO site were the abundance of genes involved in acquisition of N and S were highest in this site. This matches the ecological need within SJO due to the low nitrogen levels measured[10, 11] and likely low S conditions in this rain-fed (ombrotrophic) site. We also observed patterns in P and Fe acquisition genes across the three sites. Genes involved in the acquisition of phosphonate and organophosphorus compounds were enriched in SJO relative to QUI and BVA. Low concentrations of P have been detected at SJO [10, 11], coinciding with gene abundances at the community level where microbial populations in SJO putatively scavenge P from organic compounds as opposed to direct ion uptake, whose genes were abundant in BVA and QUI[11, 13]. Additionally, Fe uptake may potentially be mediated by siderophores in QUI and SJO. This is consistent with low concentrations of Fe measured in these soils, and the need for effective scavenging strategies for Fe [13, 63].

A defining feature of the sites in this study is the gradient of acidity correlating to decreasing nutrient concentrations (Supplementary Table 1). This is keenly reflected at the community level where we see statistically significant patterns of gene abundances associated with pH coping strategies. In BVA, the least acidic of the three sites, there is an abundance of genes involved in consuming protons via the decarboxylation of amino acids, such as arginine and glutamate. This coping strategy is commonly utilized by neutrophilic organisms[64], and could suggest the prominence of neutrophilic populations within BVA needing to cope with changes in pH from flooding events. Alternatively, in QUI and SJO, the more acidic soils, microbial communities appear to be utilizing alternative pH survival strategies, such as, ammonia generation and proton pumping which are more common in acidophilic organisms[64].

Lastly, a very notable finding is the strong signal in SJO, where genes for prokaryotic defense mechanisms were enriched. As the most nutrient limited of the three sites, it is possible that ecological competition, induced by nutrient stress, is common among SJO microbial populations[65]. The abundance of CRISPR-Cas systems in SJO suggest phages are a key component of the ecological dynamics of these soils. The use of phage infection as a mechanism to outcompete competitor populations has been shown in competing non-infected and infected populations [66, 67]. It is thus possible that the high abundance of phage immunity genes observed in SJO evidences a prevalent mechanism for competition between populations. While we cannot definitively conclude if the acquired phage immunity in SJO is utilized within populations or a stochastic property of the community; it does appear that the viral community plays a large role in these microbial communities. While microbial warfare is probable in low nutrient environments, we also recovered a high abundance of genes involved in TA systems. The frequency of persister cells, dormant cells which down regulate macromolecule biosynthesis and function towards survival, within microbial populations is linked to TA systems[68]. The infrequency of nutrient influx to SJO, an ombrotrophic rain-fed peatland, suggest that many populations enter dormant states and are overall less active than those in BVA and QUI. The prevalence of biological interactions as suggested by gene frequencies, is a growing component in microbial functioning research [69, 70] and whose role has not been previously considered in tropical peatlands.

Considering the relative abundance of genes involved in foraging mechanisms (C, N, S, P, and Fe) and the abundance of prokaryotic survival strategies between the three sites; our hypothesis that nutrient concentrations influence the functional landscape is well supported in the community scale observations. Furthermore, this supports that ecosystem level views (*i*.*e*. nutrient concentrations), at the microbial level are consistent with macroecological views that abundance and diversity of “food” can be correlated to the diversity and foraging mechanisms of the fauna that inhabit the environment[71–73].

### Differences in MAG-resolved populations mediating biogeochemical pathways across the PMFB

Our findings, at the community scale, demonstrate site-specific differences in gene abundance across the three sites, however linking these functions to microbial populations requires further exploration at a more detailed, genome-scale, gene-linkage resolution through the binning of contigs into MAGs. When comparing recovery of MAGs across peatland sites BVA was partially underrepresented; this is attributable to the high diversity of community members and requires higher sequencing coverage than was affordable in the current study. Beyond that, the frequency of MAGs recovered from the sites are consist with 16S rRNA gene abundances, as previously reported by Finn, et al, 2020. For instance, MAGs classified as *Methanomasilliicocales* were exclusively recovered from QUI and SJO, while *Methanobacteriales* from BVA. Further, MAGs classified as *Acidobacteriota* were predominately recovered from SJO consistent with the abundance of *Acidobacteriota* detected from previous 16S rRNA gene studies[13].

Our MAG analyses detail the putative microbial populations involved in C, N, and S cycling across the nutrient gradient in the PMFB. We found that *Acidobacteriae* is the most common contributor to carbon degradation within the nutrient poor sites of QUI and SJO, while the high nutrient content of BVA lead to a greater diversity of contributing members. These findings coincide with similar *Acidobacteriae* dominance found in other genome-centric studies that considered permafrost peatlands [23, 74]. This represent then an important conservation of the polysaccharide degradation function within a geographically widespread group[75], as well as, heterotrophic degradation process is distinct in peatlands by *Acidobacteriae*. Also, our study identified microbes with potential to participate in functions not previously considered. For instance, MAGs putatively capable of CH_4_ oxidation showed almost no similarities to those identified in other studies[76]. Future studies, however, should evaluate the quantitative contribution of these identified groups.

We also identified variations in the abundance of MAGs capable of carrying out biogeochemical transformations for inorganic compounds, notably those involved in NO oxidation and sulfate reduction. The flux of N_2_O were previously reported to be high at both BVA and SJO [13]. In the case of BVA, we recovered a higher diversity of *norBC* containing MAGs, even though the number of MAGs recovered from this ecosystem is lower. It has been shown that higher N_2_O flux is associated to higher gene abundance and diversity in *norBC*, which could, in part, explain the high rates in BVA[77, 78]. We noticed key differences in putative strategies of NO reduction across all sites, with many MAGs from BVA and SJO conserving energy via this reaction (*qNor*). Further, we see many NO reducers (predominately *Thermodesulfovibrionia* and UBA9217) are capable of dissimilatory sulfate reduction, indicating that some organisms can contribute to both nitrogen and sulfur cycles potentially outcompeting methanogenesis[19] and thus capable of negatively regulating it. The possible function of *Acidobacteriae* to reduce NO has been previously suggested [79] but also overlooked[75] due to the lack of experimental evidence of such activity. In sites like BVA[13, 19] NO reduction via *qNor* is likely an important pathway for the release of N_2_O from the soils and should be quantitatively evaluated.

### Conclusion

Overall, this study provides insights into the effects of geochemical variation (nutrients, pH, others) on the functional landscapes of the microbial communities found within the PMFB. Our observations of significant gene patterns associated to nutrient acquisition are illustrative of the effects that an ecosystem’s geochemistry can have on the microbial populations that inhabit these environments. Further, our results at the genome-resolved scale further suggest that local environmental conditions influence the partitioning of functions involved in C, N, and S cycling. Future efforts need to combine genome-centric data with metabolomics and biogeochemical data to better improve our ability to accurately predict soil carbon degradation and GHG emission processes from these important ecosystems in the Amazonia.

## Acknowledgements

We thank the assistance of Daniela Boullosa, Lenin Caritimari, Elda Rodrigues inf field sampling activities, as well as the assistance of Cpt. Isidoro Pacaya and other members of the Buena Vista and San Jorge communities in the Tahuayo River, Loreto, Peru. All field work and sampling has been approved under permit number

## Data Availability Statement

Metagenomes used in this study are deposited with JGI under the GOLD Biosample IDs Gb0153535 - Gb0153559. All MAGs are provided at the Figshare data repository: **10.6084/m9.figshare.21433599**. The data and scripts used in this manuscript are publicly available at: https://github.com/Hinsby/MetagenomeAmazonPeatlands2022

## Originality-Significance Statement

This work uncovers the possible functional constrains imposed by distinct geochemistry across Amazon peatlands that not only affect metabolic types (acting in the C, N, S, and others cycling) but also biological processes of microbial communities using distinct defense and cellular homeostasis mechanisms for their functioning.

Hundreds of new metagenome-assembled genomes (MAGs) for Bacteria and Archaea, are provided here to seek a more detailed evaluation of the putative contribution of extant microbes in tropical peatlands. Such analyses, for instance, showed phyla putatively capable to participate in both nitrogen and sulfur cycle, or that the novel functions suggested for Acidobacteria in northern peatlands in early steps of organic matter degradation is also present and perhaps in a more pervasive fashion in tropical peatlands. This report is the most comprehensive metagenomic effort for tropical peatlands, also is the first one for Amazon peatlands, it details multiple findings that can be used in current efforts studying the carbon sinking capacity of tropical regions, greenhouse gas emissions plus our need to better understand the biological functioning of tropical soils.

