## Supplementary Table 1 and Figure 1 for "Genes and genome-resolved metagenomics reveal the microbial functional make up of Amazon peatlands under geochemical gradients"

**Supplementary Table 1.** Summary of study site characteristics and soil properties from the Pastaza-Marañón Foreland Basin

| | **Buena Vista (BVA)** | **Quistococha (QUI)** | **San Jorge (SJO)** |
|---|---|---|---|
| Soil Category | Minerotrophic | Mixed | Ombrotrophic |
| MAT (°C) | 26 | 26.4 | 26.4 |
| MAP (mm) | 2506.8 | 2987.4 | 2506.8 |
| Bulk Density (g $cm^{-3}$) | 0.09 ± 0 | 0.11 ± 0 | 0.1 ± 0 |
| pH | 5.6 ± 0.1 | 3.7 ± 0.1 | 2.5 ± 0 |
| $CO_2$ (mean flux ug C $m^2$ $h^{-1}$) | 65.3 ± 15.4 | 29.9 ± 35.1 | 59 ± 24.7 |
| $CH_4$ (mean flux ug C $m^2$ $h^{-1}$) | 378.4 ± 636.4 | 407.1 ± 529.9 | 189.9 ± 360.6 |
| $N_2O$ (mean flux ug C $m^2$ $h^{-1}$) | 26.9 ± 15 | 72.7 ± 83.1 | 69.5 ± 69.5 |
| DOC (mg $L^{-1}$ water) | 27.68 ± 1.8[a] | 44.01 ± 31.1[b] | 55.5 ± 28.2[b] |
| DIC (mg $L^{-1}$ water) | NA | 0.44 ± 0.6 | 1.63 ± 0.7 |
| Nitrite (mg $L^{-1}$ water) | 0.04 ± 0.02[a] | 0.02 ± 0.06[b] | 0.006 ± 0.0[b] |
| Nitrate (mg $L^{-1}$ water) | 0.23 ± 0.1[a] | 2.39 ± 6.4[b] | 0.96 ± 1.4[b] |
| Ammonium (mg $L^{-1}$ water) | 0.63 ± 0.2[a] | 0.25 ± 0.2[b] | 0.24 ± 0.2[b] |
| Sulfate (mg $L^{-1}$ water) | 0.67 ± 0.3 | 1.46 ± 0.2 | 0.11 ± 0.1 |
| Phosphate (mg $L^{-1}$ water) | NA | 0.16 ± 0.2 | 0.03 ± 0 |
| Na (mg $L^{-1}$ water) | NA | 2.01 ± 0.2 | 0.31 ± 0 |
| Na (mg $kg^{-1}$ dry soil) | 273.8 ± 66 | 79.4 ± 24.7 | 39 ± 7.4 |
| Mg (mg $kg^{-1}$ dry soil) | 729 ± 71.2 | 374.6 ± 43.6 | 276.3 ± 59.5 |
| P (mg $kg^{-1}$ dry soil) | 1113.1 ± 339.6 | 1048.2 ± 27.6 | 676.5 ± 139.6 |
| S (mg $kg^{-1}$ dry soil) | 4746.7 ± 718.3 | 2786.7 ± 547.7 | 1863.3 ± 161.1 |
| K (mg $kg^{-1}$ dry soil) | 1763.3 ± 741.4 | 452.9 ± 18.4 | 372.6 ± 110.9 |
| Ca (mg $kg^{-1}$ dry soil) | 6516.1 ± 244.1 | 4875 ± 1301.1 | 952 ± 269.2 |
| Mn (mg $kg^{-1}$ dry soil) | 70.3 ± 27.7 | 49.2 ± 12.3 | 35.5 ± 7.1 |
| Fe (mg $kg^{-1}$ dry soil) | 7.1 ± 2.8 | 3.3 ± 3.8 | 1.3 ± 0.1 |
| Ni (mg $kg^{-1}$ dry soil) | 7.4 ± 2.4 | 2.3 ± 0.1 | 1.9 ± 0.3 |
| Cu (mg $kg^{-1}$ dry soil) | 9.8 ± 3.3 | 3.2 ± 0.3 | 6 ± 0.9 |
| Zn (mg $kg^{-1}$ dry soil) | 29 ± 11 | 10.6 ± 3.6 | 20.6 ± 5.5 |

*Unless noted all values are from Finn et. al., 2020. [a]Water values from Buessecker et. al. 2021. [b]Water values are the averages from Finn et. al., 2020 and Buessecker et. al. 2021. MAT = Mean Annual Temperature. MAP = Mean Annual Precipitation. DOC = Dissolved organic C. DIC = Dissolved inorganic carbon. NA = Measurements not taken.*

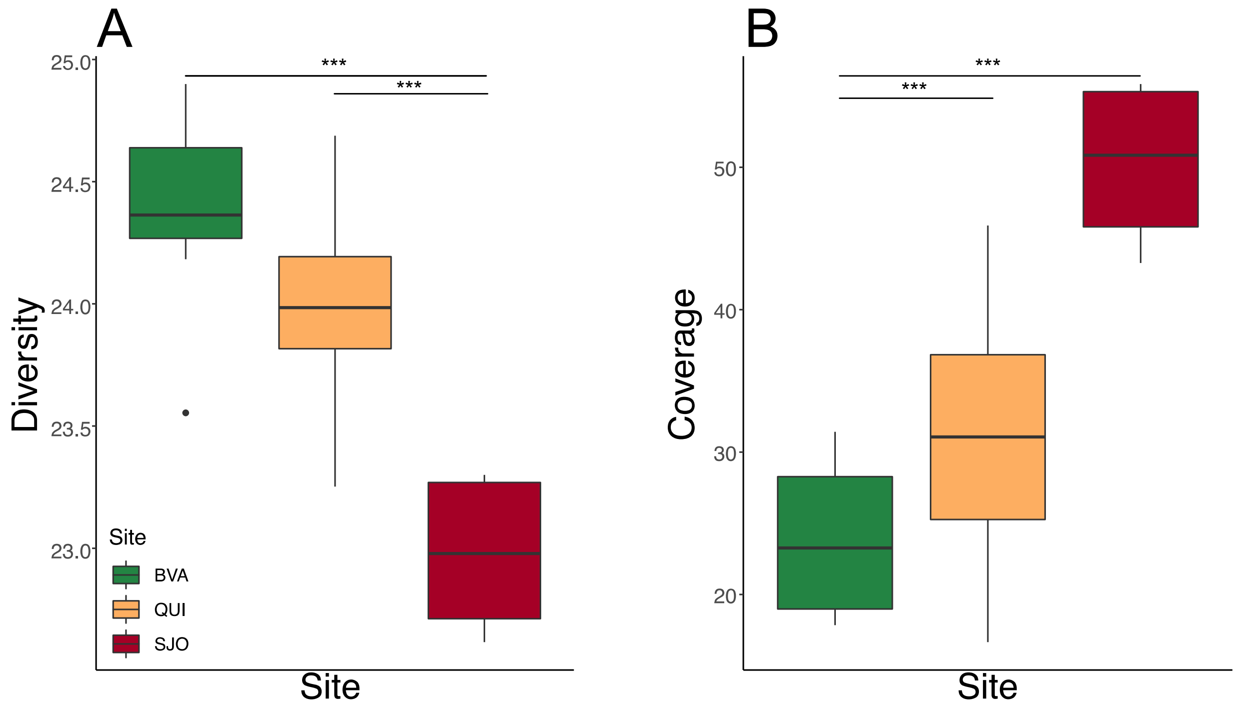


***Supplementary Figure 1.*** *Average alpha diversity (A) and coverage estimates (B) from metagenomes. Black likes above boxplots indicate statistically significant differences between those sites (two-way ANOVA, p < 0.05). BVA; Buena Vista, QUI; Quistococha, and SJO; San Jorge.*
