## Supplementary material for "Genes and genome-resolved metagenomics reveal the microbial functional make up of Amazon peatlands under geochemical gradients": Supplementary_File_Descriptions.docx

Supplementary File 1. Description of sample metagenomes and summary statistics of subsequent assembly and contig binning.

Supplementary File 2. Log2fold change of KO’s significantly different between BVA, QUI, and SJO.

Supplementary File 3. Characteristics and Abundance of Metagenome Assembled Genomes recovered from the Pastaza-Marañón Foreland Basin

Supplementary File 4. KEGG definitions used to define metabolic pathways assigned to each MAG in Figure 4.
